# Proapoptotic RHG genes and mitochondria play a key non-apoptotic role in remodelling the *Drosophila* sensory system

**DOI:** 10.1101/2021.01.15.426850

**Authors:** Amrita Mukherjee, Sinziana Pop, Shu Kondo, Darren W Williams

## Abstract

Caspases are best known for their role in programmed cell death but have also been found to be important in several non-apoptotic phenomena such as cell fate specification, cell migration and terminal differentiation. The dynamics of such sub-lethal caspase events and the molecular mechanisms regulating them are still largely unknown. As more tools for visualizing and manipulating caspase activation *in vivo* become available, greater insights into this biology are being made. Using a new and sensitive *in vivo* effector caspase probe, called SR4VH, we demonstrate that effector caspases are activated in pruning sensory neurons earlier than previously thought and that the level of caspase activation in these neurons is consistently lower than in neurons undergoing cell death. We reveal that Grim and Reaper, two of the four pro-apoptotic RHG proteins, are required for sensory neuron pruning and that disrupting the dynamics of the mitochondrial network prevents effector caspase activation in both pruning and dying sensory neurons. Overall, our findings demonstrate that a sublethal deployment of the ‘apoptotic machinery’ is critical for remodelling dendrites and also reveal a direct link between mitochondria and sensory neuron cell death *in vivo*.

## Introduction

Cysteine aspartate-specific proteases (Caspases) are key mediators of programmed cell death by apoptosis. Apoptosis is a universal mode of cellular destruction in metazoans and is critical for the development of tissue architecture and organ systems (1). Whilst the elimination of whole cells is important for sculpting tissues, it has become clear that caspases also play non-apoptotic roles including cell fate specification, migration and terminal differentiation of cell shape/function (2). In the nervous system, apoptosis plays a significant role in network construction, where as many as 50% of the neurons generated are removed (3). Caspases are also known to function non-apoptotically, during the refinement of neuronal arborizations and during synaptic plasticity (4) (5). Insects undergoing complete metamorphosis have long been a powerful *in vivo* model for studying regressive developmental phenomena (6,7). The nervous systems of such metamorphic insects are dramatically reshaped during the transition between larval and adult forms by the removal of redundant larval neurons and by the repurposing of neurons that survive, prune and then regrow to generate *de novo* adult-specific arborizations (8). Our previous work and that of others have shown that caspases are important during the remodelling of the sensory nervous system where they are activated in dying neurons and within the dendritic branches that are removed during pruning (9,10). In mammals, caspases and inhibitors of apoptosis have been shown to be critical for trophic factor mediated axon fragmentation, in both sensory and sympathetic neurons (11-13) (for review see (14)).

In mammals, we know that caspase activation during cell death can be initiated by one of two pathways: the ‘intrinsic’ mitochondrial pathway and the ‘extrinsic’ cell death receptor pathway. The majority of studies in neurons have focused on the intrinsic ‘mitochondrial’ pathway where caspases are present in cells as proenzymes and are activated in a hierarchical manner. In mammalian cells, cytochrome c, released from the inner mitochondrial membrane, forms a complex with Apaf-1 and an initiator caspase, Caspase-9, which then allows the self-activation of Caspase-9. Active Caspase-9 then cleaves and activates effector caspases including Caspase-3, which in turn targets a large number of cellular proteins (see (15,16) for reviews). The release of cytochrome c is essential in mammals but appears to be dispensable for the formation of the apoptosome and cell death in *Drosophila* (17). In flies, the initiator caspase DRONC is activated by the Apaf1 homolog, Ark (18), which then cleaves the effector caspases Drice and Dcp-1. These are ultimately responsible for executing almost all of developmental cell death in flies (19). Key proapoptotic regulators in flies are Reaper, Hid, Grim and Sickle. These ‘RHG proteins’ remove the inhibitor of apoptosis proteins (IAP) which normally bind to and degrade DRONC (for review see (20)). These IAP antagonists, have analogues in mammals, Smac/Diablo and Omi/Htr2A, which are intimately associated with mitochondria (21) (22). When RHG proteins are expressed in mammalian or *Drosophila* cells, they localise to the mitochondria and this is required for their pro-apoptotic function (23-26). Mitochondrial localisation of Reaper, Grim and Sickle depends on the presence of a GH3 domain (24,26) whereas Hid requires a mitochondrial target sequence and the Cyclin-dependent kinase 7 (Cdk7) protein (27,28). To localise to mitochondrial membrane, Reaper can either interact directly with the lipids in the outer mitochondrial membrane via its GH3 domain (29) or form a multimeric complex with Hid (30) that contributes to autoubiquitination and degradation of DIAPs (29). Although DRONC and Drice also localise to the mitochondria in cultured *Drosophila* cells (31), where caspases localise *in vivo* within *Drosophila* neurons, is still an open question.

Although the role of the involvement of cytochrome c and an intrinsic pathway of caspase activation in *Drosophila* has remained controversial there is a growing body of evidence to suggest that mitochondria play a key role in caspase activation in dying cells (17). Mitochondria act as critical nodes for signal integration within cells, with their structure and dynamics also being directly related to caspase activation (32,33), but whether they play a role in non-apoptotic caspase function is largely unexplored.

In this paper we reveal the dynamics of caspases activation and the role of mitochondria in the restructuring of the sensory nervous system during *Drosophila* metamorphosis. We show that effector caspases are activated much earlier than previously known in the dorsal dendritic arborization C (ddaC) neurons as they undergo dendrite pruning during metamorphosis. We also find that caspase activation is at a substantially lower level in these pruning neurons than in dendritic arborisation (da) neurons that die. We reveal that two of the proapoptotic RHG genes, Grim and Reaper, are required for sensory neuron pruning and show that mitochondria play a key role in dendrite remodelling. We also uncover a direct link between mitochondrial function and caspase activation during neuronal cell death in sensory neurons in *Drosophila.*

## Materials and Methods

### Fly stocks

The following fly lines were used: *UAS-CD8::PARP::Venus* (10)*, UAS-Mito::GFP* (BL 8442), *ppk-GAL1.9* (expressed in Class IV neuron (34)), *ppk-GAL4* (BL32078 and BL32079) (expressed in class IV and class III neurons), *19-12GAL4 UAS-CD8::GFP* (35), *UAS-RHG miRNA* (36), *UAS-TFAM* (37), *UAS-Mito::XhoI* (38), *UAS-Milton RNAi* (BL44477), *UAS-Miro RNAi* (BL51646), *UAS-Marf RNAi* (BL 55189), *UAS-Opa1 like RNAi* (BL32358), *UAS-Drp1 RNAi* (BL27682), *UAS-Dicer2* (BL 24648), *H99* deficiency (BL 1576), *XR38* deficiency (BL 83151), *rpr^SK3^*/TM6B (This study), *hid ^SK6^*/TM6B (This study), *grim^A6C^* (BL32061), *UAS-Drp1WT* (39), *elav-GAL4 ^C155^* (BL 458), *UAS-RedStinger* (BL 8545), *SOP-FLP* on X (a gift from Tadashi Uemura), *nSyb-GAL4* (BL51635), *UAS-SR4VH* (40). For generating the modified mosaic clones with a repressible cell marker (MARCM) experiment for *H99*, similar procedures were followed as described previously(41).

### Immunohistochemistry and imaging

Larvae and pre-pupae were dissected as described previously (42). The fillet preps were fixed in freshly prepared 4% formaldehyde for 20 minutes at room temperature. The fixative was washed off with PBST (0.3% TritonX-100). The preps were then blocked in 5% BSA in PBST for 1h at room temperature and incubated in appropriate mix of primary antibodies overnight at 4°C. The primary antibody solution was washed off the next day and samples incubated in secondary antibody solution overnight at 4°C. After washing in PBST and then PBS the following day, the fillet preps were mounted on poly-L-lysine coated coverslips. The samples were serially dehydrated through an ethanol series, washed twice in xylene and mounted in DPX.

The following primary antibodies were used: Mouse anti-GFP (1:400, Abcam), Rabbit anti-PARP (1:500, Abcam ab2317), Mouse anti-EcR (1:5, DSHB), Guinea pig anti-Sox14 (1:500, gift from Fengwei Yu), Rabbit anti Dcp-1 cleaved (1:100, Cell Signalling). All secondary antibodies were used at 1:500 dilutions, obtained from Jackson Laboratories.

For live imaging the pre-pupae were mounted under a coverslip on a standard glass slide. A very small amount of Halocarbon oil (Voltalef) was added to the contact point between the sample and the coverslip and pressed down lightly onto four small 2 mm balls of dental wax to act as spacers. This preparation was then imaged immediately on the microscope. In order to image pupae, white pre-pupae were selected and aged at 25°C for the required duration in a humid chamber and dissected out of the pupal case before mounting in the same way as pre-pupae. We used Zeiss LSM 510 or LSM 800 and Plan-Apochromat 40x/1.3 objective for imaging and the Olympus FV3000 scanning inverted confocal system run by FV-OSR software using a 60X 1.4NA silicon immersion lens (UPLSAPO60xSilcon).

To obtain intensity measurements of neurons expressing SR4VH we used Line plots; the Plot Profile tool in Fiji was used to extract raw fluorescence intensity values for the RFP and Venus channels. The values were then imported into MATLAB (R2018a, MathWorks) and normalised by dividing all fluorescence intensity values to the maximum value for the RFP channel encountered along each Line at each timepoint such that all fluorescence intensity along Line plots have a common scale from 0 to 1, with 1 being the highest value encountered in the RFP channel along that Line and at that timepoint.

Dying SR4VH cells in the wing pouch were counted from one optical slice taken from the middle of the Z-stack and analysed using the Kruskal-Wallis test to compare mean ranks, as the data failed to meet the normality and homogeneity of variances assumptions of one-way ANOVA. Statistically significant findings were followed up with pairwise Mann-Whitney tests, with p values adjusted using a Bonferroni correction (p values were multiplied by the total number of pairwise tests performed for each multiple comparison).

To measure the number of mitochondria, we used ImageJ Multi-point tool to count the number of mitochondria and ImageJ segmented line tool to measure the total length of dendrites and calculated the number of mitochondria per 100 microns of dendrite length. To compare several genotypes to control, we performed the Kruskal-Wallis test followed by Dunn’s test, with p values adjusted using a Bonferroni correction.

We used ordinary one-way ANOVA for statistical significance. For analysing pruning phenotypes, we divided the phenotypes in three categories: firstly- no phenotype, these appear like wildtype i.e. field imaged is completely clear of dendrites; second - clearance defect, where the main dendrites are separated from the cell body but not cleared from field; third, severing plus clearance defects, where the primary dendrites remained attached to the cell body and cut dendrites in the vicinity are not cleared. For each genotype we imaged 2 – 3 abdominal neurons per animal. N numbers represent total number of neurons imaged.

### CRISPR mutagenesis

New null alleles of *rpr* and *hid* were generated using the transgenic CRISPR system as described (43). The target-specific 20-bp sequences of the gRNAs are as follows: *rpr*: GGCATTCTACATACCCGATC, *hid*: TGAACTCGACGCTACGTCAT. We screened candidate mutant lines by Sanger sequencing and selected those that carry a frameshift-causing indel mutation in the respective genes. The molecular lesions of the new alleles are as follows: *rpr^SK3^*: CTACATACCC-ATCAGGCGAC, *hid^SK6^*: GCGCCGATGA-----------GTTCATCGGG, where deleted bases are indicated as dashes.

## Results

### Visualizing caspase activation within sensory neurons at the onset of metamorphosis

The larval sensory system of *Drosophila melanogaster* has bilateral, segmentally repeated clusters of neurons within the dorsal abdominal body wall. Each of these clusters contain thirteen sensory neurons, that are uniquely identifiable, six of these neurons are dendritic arborisation (da) sensory neurons which have characteristic tree-like, peripheral arborisations (44) (Fig.1A-C). In insects, like other arthropods, the cell bodies of sensory neurons are located in the periphery and axons from them track through peripheral nerves to terminate in the central nervous system (CNS). (Fig. 1A)

**Figure 1:**
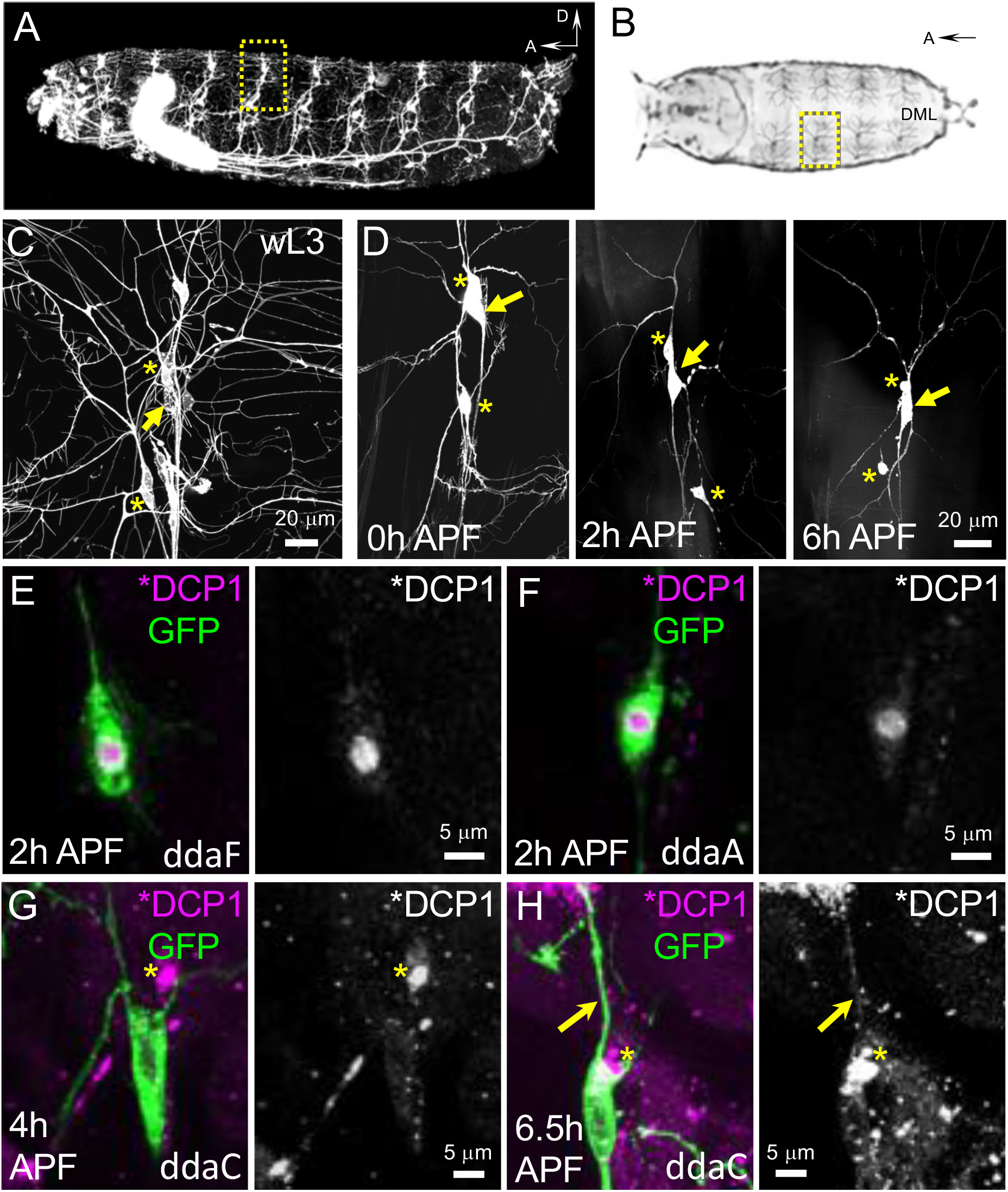
The sensory system of *Drosophila* undergoes extensive remodelling at metamorphosis. (**A**) A 2^nd^ instar *Drosophila* larva expressing GFP in cholinergic neurons (ChaGAL4>UASGFP) shows the organisation of the central and peripheral nervous system. Segmentally repeated arrays of sensory neurons send their axons through peripheral nerves to the ventral nerve cord. D=dorsal, A=anterior. Yellow box indicates the position of the dorsal cluster of sensory neurons on the body wall. (**B**) Drawing of an early *Drosophila* pre-pupa from above with the dorsal cluster of sensory neurons indicated by a yellow box. DML = dorsal midline. (**C**) Higher magnification of the sensory neurons in a dorsal cluster, labelled by ChaGAL4>UAS CD8::GFP, the cell bodies of pruning Class IV ddaC (arrow) and dying class III ddaF and ddaA (asterisks). (**D**) Two class III neurons - ddaA and ddaF (asterisks) and one class IV ddaC neuron (arrow), imaged *in vivo* show neurons undergoing cell death and remodelling respectively at the onset of metamorphosis. These three neurons are revealed with two copies of ppkGAL4>UAS CD8::GFP (**E,F**) Class III neurons are labelled using 19-12 GAL4>UAS CD8::GFP. **(E)** Left panels show ddaF fixed and immunostained for GFP (green) and active effector caspase DCP-1 (magenta). Right panels showing immunoreactivity of active DCP-1 alone in greyscale. **(F)** shows ddaA. Strong active DCP-1 staining is seen in the nuclear region of both dying cells (**G,H**) Left panels showing ddaC labelled using ppkCD4tdGFP, fixed and immunostained for GFP (green) and active DCP-1 (magenta) right panels showing immunoreactivity of active DCP-1 alone (greyscale). Weak active effector caspase expression in the cell body and dendrites at 4h APF and active caspases can be detected in a dendrite still attached to the cell body (arrow) at 6h APF. In contrast to dying neurons, no active caspase is present in the nucleus. Asterisks mark DCP-1 nuclear staining of apoptotic sensory neurons in the vicinity.

At the beginning of metamorphosis, three of the da neurons in the dorsal cluster, ddaA, ddaF and ddaB, undergo programmed cell death (Fig.1C and D, images only refer to dying neurons ddaA and ddaF, asterisks) whereas the other three, ddaD, ddaE and ddaC, survive and are remodelled (Fig. 1 C and D, ddaC indicated by arrow) (45,46). At the onset of metamorphosis da neurons remove their larval-specific dendrites by pruning (45), migrate up the body wall and then elaborate *de novo* adult-specific arborizations (47-49).

At pupariation the dorsal cluster of neurons can be easily observed through the dorsal puparial case (Fig. 1B). By 2h after puparium formation (APF) the cell bodies and proximal dendrites of the dying neurons, ddaA and ddaF, show clear signs of disintegration (Fig.1D, asterisks). By 6h APF their cell bodies appear condensed and their dendrites have fragmented, both features being characteristic of apoptotic cells. The dendritic fragments and dying condensed cell bodies are rapidly cleared by macrophages (45). The pruning neuron, a class IV da called ddaC, shows morphological changes by 6h APF (Fig.1D, arrow) with its proximal dendrites beginning to thin and generate varicosities along the length (Fig.1D). This period of thinning and beading is quickly followed by branch severing events, after which detached dendrites undergo fragmentation and are engulfed by macrophages and epidermal cells in the vicinity (45) (50).

Our previous work using the genetically encoded effector caspase reporter, CD8::PARP::Venus, revealed that caspases are activated within ddaC neurons during pruning but only in dendritic branches after they had been cut from the cell body (10). To explore the timing of the onset of caspase activity we used an antibody that recognises the cleaved form of Dcp-1 and Drice (51). Because Dcp-1 and Drice are direct substrates of DRONC, this antibody should be able to detect caspase activation prior to the cleavage of the CD8::PARP::Venus reporter by active effector caspases. Looking at the doomed/dying neurons ddaF and ddaA, we saw that Dcp-1/Drice are robustly cleaved within each cell, with strong nuclear staining and a weaker cytoplasmic staining (Fig.1E&F). In the pruning neuron ddaC, we found a low level of staining for cleaved Dcp-1/Drice in the cell body and also within the intact proximal branches of ddaC at 6.5h APF (Fig.1 G&H, arrow). The cleaved Dcp-1/Drice immunoreactivity within the dying da sensory neuron always appeared stronger (Fig.1 E&F) than in ddaC, the pruning neuron (asterisks indicates dying sensory neurons beside the ddaC neuron Fig 1 G,H). These data, in contrast to our previous findings, indicate that caspases are active in pruning ddaC neurons early in pupariation, before dendrite branch severing has taken place. (Fig.1 H).

### SR4VH a genetically encoded probe for visualising sublethal levels of effector caspase activation

It could be that there is no effector caspase activity in pruning neurons at this earlier time point as a number of studies looking at non-apoptotic developmental events, e.g arista morphogenesis (52) and border cell migration (53), found that only initiator caspase (DRONC) activity is required. Encouraged by our initial observations with the DCP1 stainings (Fig. 1) and to interrogate this idea further, we utilized our newly developed genetically encoded probe to gain further insight into the timing of effector caspase activity and how this relates to changes in the structure of the dendrites of pruning neurons. We have previously demonstrated SR4VH to be sensitive and capable of detecting the temporal details of caspase activity in newly born postembryonic neurons that undergo hemilineage-specific patterns of cell death (40). The probe has an mRFP1 red fluorescent protein fused together with the Src64B myristoylation signal and a Venus fluorescent protein with a histone H2B nuclear localisation signal (Venus::H2B). These two fluorescent domains are joined by a linker containing four repeats of the DEVD sequence, an optimal effector caspase cleavage site shown to be effectively cleaved by the fly effector caspases, Drice and Dcp-1 (54). We call this new probe ‘*SR4VH*’ because of its structure (***S**RC::**R**FP::**4**xDEVD::**V**enus::**H**2B*) (Fig.2A). Although similar in design to the previously published *Apoliner* probe (55), SR4VH is different in that it contains four tandem DEVD sequences, not just the single caspase cleavage site from DIAP1, to improve its cleavage efficiency. It also uses a different localisation signal to tether the reporter to the membrane compartment (40). With *Apoliner* we found that large amounts of newly generated reporter protein accumulate in the Golgi apparatus within the cell bodies of the sensory neurons, often obscuring the nuclear signal (Supplemental Fig. 1C, D).

**Figure 2:**
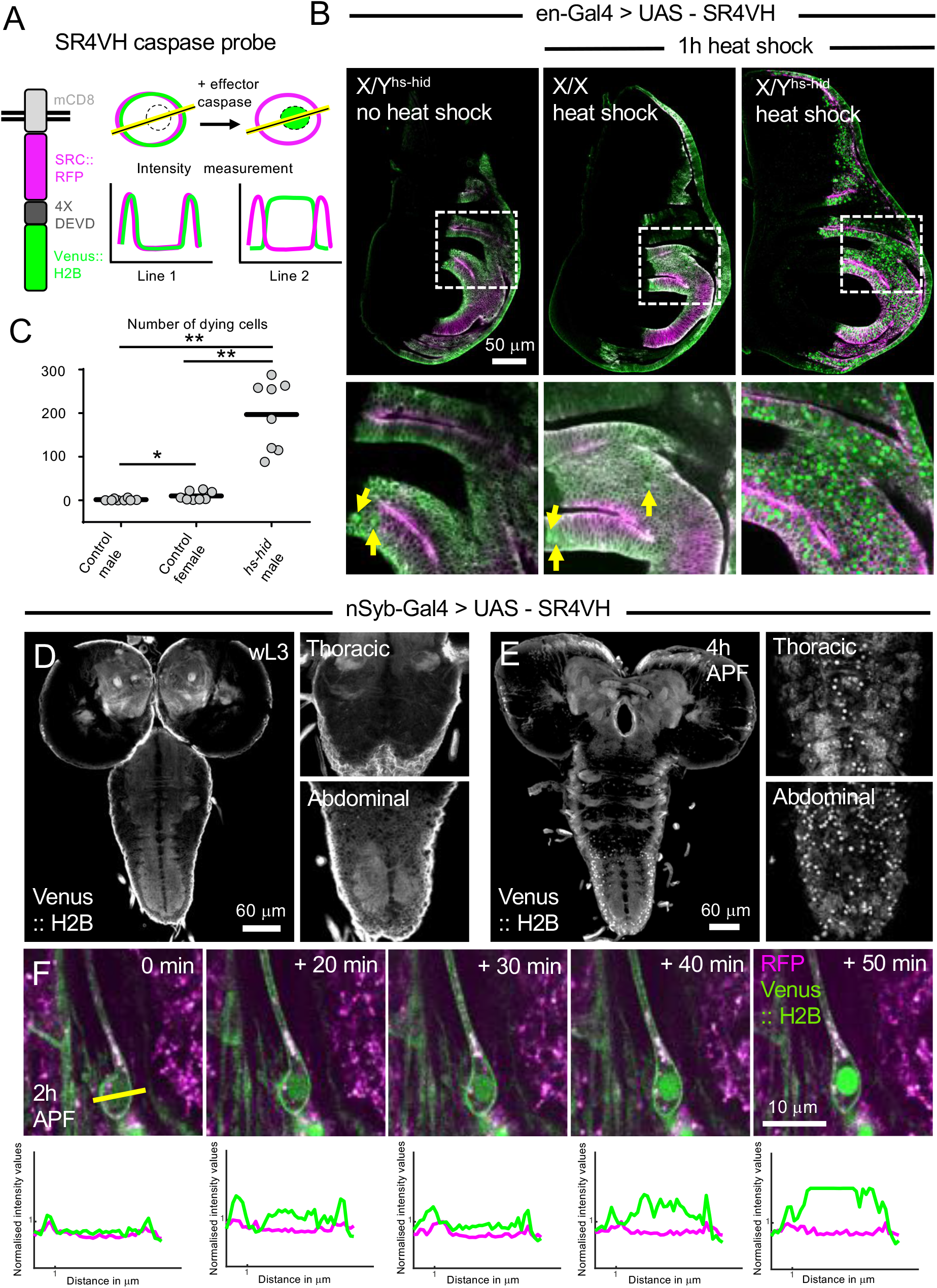
The genetically encoded effector caspase reporter SR4VH reveals cell deaths *in vivo*. **(A)** A schematic of SR4VH showing the reporter changing location following cleavage and how intensity measurements across the neuronal cell body provide a readout of caspase activity. (**B**) Wing discs from 3^rd^ instar larvae expressing SR4VH under the control en-GAL4, GFP (green) and RFP (magenta). Left panel showing a disc from a “no heat shock” control male X/Y^hs-hid^ and middle panel showing disc from “heat shocked” control female X/X containing no hs-hid transgene. Both control conditions reveal a few cells with nuclear GFP signal (arrows). Right panel shows a wing disc from a “heat shocked” male X/Y^hs-hid^, containing many cells with nuclear localised GFP. (**C**) Quantification of the number of dying cells in the wing discs in B. (**D**, **E**) nSyb-GAL4>UAS-SR4VH expressing larval and pre-pupal nervous systems dissected, fixed and immune-stained for GFP (grey). Values are reported as mean ± standard deviation and p values are reported as * for p<0.05 and ** for p <0.01. **(D)** Left panel showing whole CNS from 3^rd^ instar larva and right panels shows magnified images of the thoracic and abdominal regions of the VNC with no nuclear GFP signal. **(E)** Left panel with whole CNS from prepupae 4 hours after puparium formation (APF) along with magnified images of the thoracic and abdominal regions showing cells with nuclear localised GFP. The abdominal region has many neurons undergoing hormonally induced cell death (right hand panels). (**F**) Upper panel shows sequence of stills from a timelapse movie of dmd1 neuron in a pre-pupa expressing SR4VH under the control of elav^C155^GAL4. Imaging starts at 2h APF. The cleaved GFP accumulates in the nuclei of the dying dmd1 neuron over the time course. Lower panel shows the normalised fluorescence intensity plots of dmd1 neuron at each of the time points displayed.

To validate the SR4VH reporter we imaged it within the developing wing imaginal discs, a tissue where sporadic developmental apoptosis has been well characterised (56). When expressing SR4VH in the posterior compartment of wildtype wing discs, we found small clusters of cells with clear nuclear localised Venus (Fig.2B, yellow arrows). We tested the status of activated effector caspases by using the cleaved Dcp-1 antibody within these clusters and found nuclear localised Venus expressing cells colocalised with Dcp-1 immunoreactivity. The cleaved Dcp-1 signal within these cells independently confirms that the clusters with nuclear localised Venus have active caspases and are undergoing apoptosis (Supp. Fig.1A).

To determine whether SR4VH could report on a rapid induction of cell death we activated the apoptotic pathway by driving the expression of the proapoptotic gene *head involution defective (hid).* Using a heat shock promoter based *hid* construct (*hs-hid*) on the Y chromosome, we shifted larvae to 37°C for 1 hour then fixed, processed and imaged the tissue (Fig.2B). After an hour incubation at 37°C, we found a significantly higher number of cells with nuclear localised Venus in *hs-hid* males, compared with the same genotype kept at 22°C and also compared to heat-treated females, that do not carry the *hs-hid* transgene. Although our heat shock treatment may have had a small effect i.e. a few additional cells are dying in Control females (heat shocked but not carrying the *hs-hid* Y chromosome) than Control males (not heat shocked carrying the Y *hs-hid* chromosome), the numbers of cells dying as reported by SR4VH expression were far greater in the heat shocked males carrying the *hs-hid* gene than either type of Controls (Fig.2B&C). We see that when SR4VH is cleaved, the Venus fragment accumulates in the nucleus. These data reveal that SR4VH can be used to accurately report on both normally occurring apoptotic deaths in developing imaginal discs and also when the apoptotic pathway is rapidly induced experimentally within the same tissue.

To see how SR4VH reports on apoptosis within the central nervous system (CNS), we imaged it before and after the onset of metamorphosis. When expressing SR4VH under the control of *nSyb-GAL4*, the neural synaptobrevin driver, we found the cell membranes of fully differentiated neurons were evenly labelled and they showed no cleavage of the probe throughout larval life (Fig.2D). We then looked at this same genotype four hours after the onset of metamorphosis (Fig.2E) and found a large number of neuronal cell bodies, with nuclear localised Venus (compare Fig.2D&E). These cell deaths occur throughout the nervous system but the largest number of neurons with nuclear Venus signals occur in the abdominal neuromeres (the region that undergoes the most dramatic remodelling at metamorphosis). These cells with nuclear Venus at 4h APF are among the class of neurons that are known to undergo hormonally-gated programmed cell death (8,57).

### Using SR4VH to visualise effector caspase activity live, in single neurons

To obtain insights into the detailed timing of caspase activation we wanted to monitor the dynamics of caspase activation ‘live’ in single cells, in intact animals. We initially focused on imaging SR4VH in the dorsal multiple dendrite neuron 1 (dmd1) in pre-pupae as it is easily identifiable, its cell body and neurites show strong immunoreactivity for cleaved Dcp-1/Drice (data not shown) and is known to be rapidly removed during early metamorphosis. We imaged dmd1 once every 10 mins through the puparial case starting from 2h APF (Fig. 2F and Supp movie 1). We saw the accumulation of Venus in the nucleus over a span of 30 minutes. Measurements of the separate Venus and RFP channels reveal the progression of the Venus marker from being in the same compartments to being spatially separated. This demonstrates that the SR4VH probe can directly report on the dynamics of effector caspase function live in a single neuron undergoing programmed cell death within an intact animal.

### SR4VH reveals early and low-level effector caspase activation within pruning neurons at the onset of metamorphosis

To image caspase activation within da neurons at the onset of metamorphosis we used two copies of pickpocket GAL4 (*ppkGAL4*) and UAS-SR4VH. We used this combination because it allowed us to cleanly visualise both the ‘doomed’ class III dorsal da neurons (ddaA and ddaF that die) and the ‘surviving’ class IV da neurons (ddaC that undergoes pruning) (Fig.3A) simultaneously. In the dying neurons, ddaA and ddaF, we found that within 20 minutes APF there is an indication of nuclear entry of Venus, and by 40 minutes APF clear nuclear accumulation (Fig.3B and Supp movie 2). This relatively rapid and robust nuclear localisation of Venus mirrors that seen for dmd1 above (Fig.2F). From the same time-lapse sequence we see that the class IV neuron, ddaC, has a lower but sustained accumulation of Venus within its nucleus from 20 minutes APF (Fig.3C,D). This nuclear entry is prior to any of the primary dendritic branches being cut (Fig. 3D). We never saw the nuclear entry of Venus in class III or class IV neurons during larval stages (Supp Fig. 1E,E’). This marks the first time that we observe a measurable change at these early stages. Previously, using the CD8::PARP::Venus probe (Fig.3E) in ddaC neurons (10), we observed very low levels of cleaved PARP (cPARP) but could not ascertain if this was background as it was barely above the levels of immunoreactivity in third instar ddaC (unpublished observations). In contrast, the cPARP-IR signal was always high in the severed branches of the pruning neurons detected between 4 – 8 h APF ((10)and Fig.3G) and in dying neurons (Fig.3F). These data from SR4VH show that caspase activation occurs very soon after the onset of metamorphosis. In contrast, we found that *Apoliner*, another published and well characterised live probe (55), did not allow us to describe these earlier events with clarity (see Supplemental Figure 1C-D).

**Figure 3:**
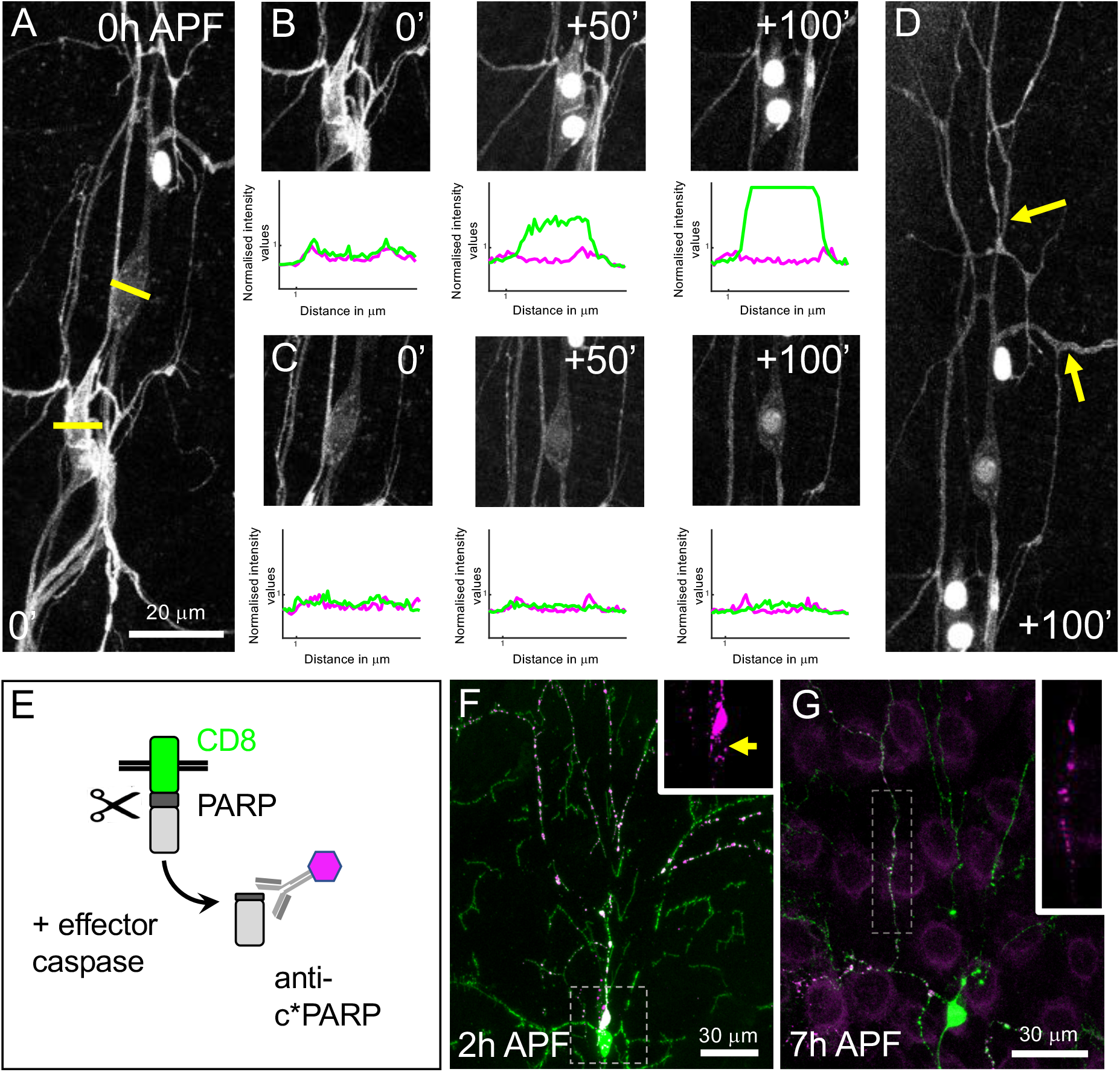
The caspase effector reporter SR4VH reveals caspase activity live in pruning and dying sensory neurons during metamorphic remodelling. **(A)** Class III and Class IV neurons were labelled using two copies of ppkGAL4 driving UAS-SR4VH. The yellow lines mark the site for sampling of intensity measurements. (**B**) Top panels show individual timepoints of the Class III neuron ddaA at which the intensity measurements were made. Within 50 minutes this neuron shows a robust accumulation of nuclear GFP signal. Bottom panel depicts the normalised intensity values for Venus and RFP plotted for the same neuron over time. (**C**) Top panel showing snap shots of the Class IV neuron at the time points at which the intensity measurements were made. There is clear but weak nuclear GFP accumulation even after 100 minutes of imaging. Bottom panel depicts the normalised intensity values plotted for the same neuron over time. (**D**) At the time point 100’+, when nuclear GFP is detected in the Class IV ddaC neuron, the dendritic branches are still intact (arrows). (**E**) Schematic of the effector caspase CD8::PARP::Venus probe that can be detected in fixed samples by immunostaining against cleaved PARP. (**F**) In a prepupa expressing 19-12 GAL4 and ppkGAL4>UAS CD8::PARP::Venus, the dying neurons (Class III) show cleaved PARP throughout the whole neuron in the cell body and dendrites. In the remodelling class IV neuron there is no cleaved PARP staining, apart for a few bright dots within the cell body at 2h APF (see arrow in inset). (**G**) In pruning ddaC neurons expressing UAS-CD8::PARP::Venus cleaved PARP immunoreactivity is evident in the branches at 7h APF.

To determine whether the differences in the levels of nuclear localization of Venus, between different cells were in fact due to differences in levels of caspase activation, we compared the ratios of RFP and Venus fluorescent channels. Using this ratiometric approach we see that the levels of the cleaved nuclear localised Venus were consistently higher in the dying neurons than the pruning neurons (Fig. 3B&C). Thus in this sub-lethal non-apoptotic context of pruning, effector caspases have a lower activity.

Our new dual colour live caspase probe (SR4VH) appears to accurately report cell death and reveals that non-apoptotic activation of effector caspases in pruning da neurons occur very soon after the onset of metamorphosis. We found a ~5 fold change in nuclear GFP signal in dying neurons compared to a 1.3 fold change in pruning neurons within the same time period, and this for the first time, gives us a quantitative readout of caspase activation live. This reveals that caspase activation is early and low in pruning neurons and occurs prior to any of the primary dendritic branches being severed (Fig. 3D arrows).

### The proapoptotic proteins Reaper and Grim are required for da neuron pruning

In *Drosophila* a major control point for apoptosis is through the post-translational regulation of the inhibitor of apoptosis proteins (DIAPs) (58). DIAP binds to both initiator and effector caspases, ubiquitylates them to target them for destruction, thus preventing apoptosis from taking place. To counter this inhibition, the proapoptotic RHG proteins (Reaper, Hid, Grim and Sickle) bind to DIAP and inhibit it. Previous work has shown that an upregulation of DIAP or a Gain of function DIAP allele in which ubiquitylation cannot take place, results in a disruption of da neuron pruning (9,10). Whilst the RHG proteins are widely known as key executors of programmed cell death during development and following DNA damage, it is not known if they play a role in regulating non-apoptotic, sub-lethal caspase function.

To test the requirement of the RHG proteins during pruning, we first blocked their function by cell autonomously knocking them down using UAS-miRNA transgene that targets *reaper*, *hid* and *grim* simultaneously (hereafter referred to as UAS-RHG miRNA) (36). By expressing these shRNAs with *ppk-GAL4* we found that pruning in the class IV neuron ddaC was disrupted (Fig. 4A and B).

**Figure 4:**
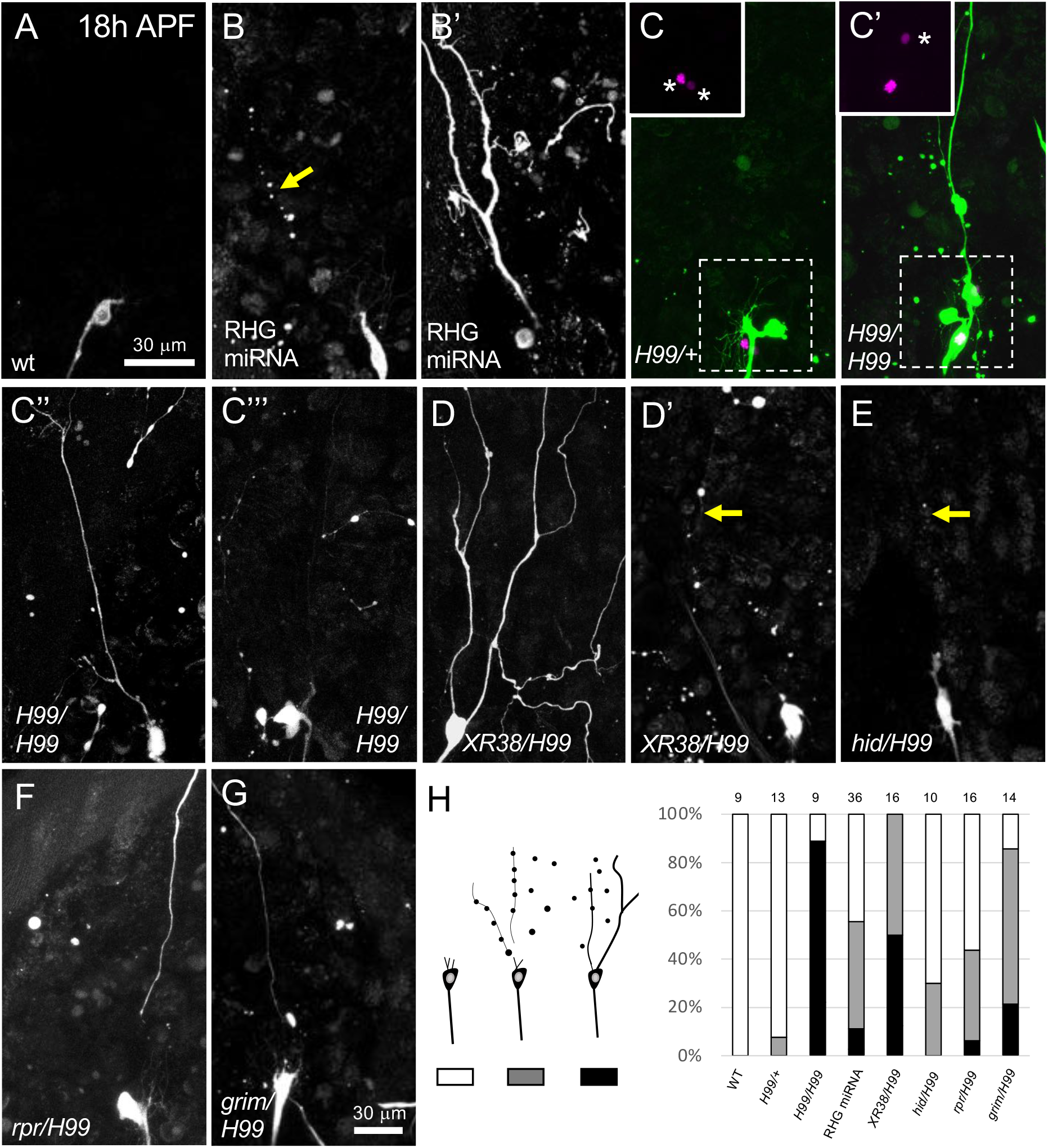
The propapoptotic RHG genes reaper and grim are important for dendrite pruning in the ddaC. All ddaC neurons were imaged at 18 h APF. (**A**) A wild-type neuron labelled using ppkGAL4>UAS-CD8::GFP shows cell body and axon with complete clearance of dendrites at 18h APF. (**B B’**) When expressing ppkGAL4>UAS-CD8::GFP, UAS-RHG RNAi to knockdown reaper, hid and grim, ddaC neurons show both severing and clearance defects (arrow). (**C**) example of control ddaC heterozygous for H99, soma has no magenta nucleus. (**C’-C’’’**) Examples of H99 homozygous MARCM neurons (magenta nucleus) in a pupa otherwise heterozygous for H99, ddaC neurons labelled using ppk-eGFP show strong severing and clearance defects, GAL80 minus neurons express RedStinger (magenta nucleus). (**D D’**) In ppkGAL4 UAS-CD8::GFP pupae with the XR38 deficiency over the H99 deficiency ddaC neurons showed strong severing and clearance defects. (**E**) Mild clearance defects in pupae deficient in hid but heterozygous for other cell death genes. (**F**) Severing and clearance defects in pupae deficient in rpr but heterozygous for the other RHG genes. (**G**) Pupae deficient in grim and heterozygous for other cell death genes, have severing and clearance defects. (**H**) Cartoon representation of the categories used when scoring for phenotypes of ddaC neuron, white - wild type, grey - clearance phenotype and black - severing and clearance. The right panel shows the percentage representation of the different categories of phenotypes as when the RHG genes are perturbed. n numbers are depicted on the top of the bars for each genotype.

We found that ddaC expressing UAS-RHG miRNA showed varying degrees of disruption, from neurons with their dendrites were completely removed (Fig. 4A), to ones with clearance defects (intact severed and fragmenting branches), to others with severing defects, that had intact primary dendrites and portions of their arborizations still attached (Fig. 4B). We found that the UAS-RHG miRNA suppresses cell death in class III da neurons (Supp Figure 2G,H) hence the variability in the pruning phenotype in ddaC we observe may be due to ppk-GAL4 driver not being strong enough to knock down all RHG proteins.

To confirm the role of the RHG proteins we looked at pruning in da neurons with the *H99* deficiency chromosome, that removes three of the four RHG genes - *hid*, *grim* and *reaper*. As *H99* homozygotes are embryonic lethal, we generated single cell mutant clones using a modified version of the MARCM (Mosaic Analysis with a Repressible Cell Marker) technique we developed previously (41). This approach allows us to see the morphology of both the single homozygous mutant clones and heterozygous control neurons side by side, in the same animal. We found that 90% of Class IV sensory neuron *H99* MARCM clones, where each clone was obtained in separate individual pupae, demonstrated a strong block in dendrite pruning. By contrast, 100% of heterozygous, *GAL80^+^* control neurons in neighbouring segments underwent pruning like wildtype neurons (Fig. 4C-C’’’ & H).

To narrow down which of the RHG genes are required for da neuron pruning, we used the chromosomal deficiency combination of *H99/XR38*. *XR38* removes the whole of the *reaper* open reading frame, some of the *cis*-regulatory region around *grim,* has a point mutation within *grim* itself and removes *sickle*. This combination results in a clear blockade of programmed cell death in the larval and adult nervous system (59). We found that this combination blocked dendrite pruning in ddaC neurons resulting in a range of phenotypes from robust branch severing to branch clearance (Fig. 4D and 4D’). In 100% of the cases we observed that pruning was disrupted, this included both branch clearance and dendrite severing phenotypes.

Following this, we then tested individual alleles of *reaper* (*rpr ^SK3^*), *hid* (*hid^SK6^*) and *grim* (*grim^A6C^*) mutants over the *H99* deficiency. With these we found that loss of both *reaper* and *grim* resulted in a disruption of pruning but, to our surprise, *hid* did not result in any suppression of branch severing and clearance was disrupted only in a small number of cases, with small fragments of dendrite remaining (Fig. 4 E – H). Since the *H99* MARCM clones, with disrupted *hid*, *grim* and *reaper* (but not *sickle*), showed very strong pruning defects with a block in dendrite severing in 90% of the cases, we chose not to investigate the role of *sickle* further, although, it is still possible that *sickle* plays a minor role in the pruning of these neurons. Out of the three RHG genes in the H99 region we found that the loss of Reaper or Grim alone resulted in weaker pruning defects than when both are removed (Fig. 4). Taken together these data suggest that Reaper and Grim play a non-apoptotic role in da neuron pruning.

### Mitochondrial physiology and transport are important for da neuron pruning

Because Reaper and Grim both have GH3 domains and are thought to interact with and degrade IAPs when localised to the mitochondria (24,26,29,30) we wanted to establish whether mitochondria play a role during the remodelling of larval sensory neurons.

To determine the requirement of mitochondria during pruning we disrupted different aspects of mitochondrial biology in single remodelling neurons. It has previously been shown that overexpression of the *Drosophila* mitochondrial transcription factor A (TFAM), dysregulates mtDNA-encoded gene expression (37). TFAM normally binds mitochondrial DNA (mtDNA) and when overexpressed in ddaC neurons, we found that dendrite pruning is disrupted. TFAM prevents both dendritic branch severing and clearance from taking place normally (Fig. 5A, B and J). As an alternative method to directly inhibit mitochondrial gene expression we expressed a mitochondrially targeted restriction enzyme MitoXhoI which is transported into mitochondria, where it cuts at a single site in the mitochondrial genome (cytochrome c oxidase subunit I) (38). We see that in ddaC neurons, expressing MitoXhoI, branch severing and clearance are significantly disrupted. (Fig. 5A, C and J). As these branch severing phenotypes resembled those seen when Ecdysone signalling is disrupted during pruning (45), we used known downstream markers to determine if there was a global impact on hormonally-gated development. We found no change in the timing or levels of EcR or Sox14 expression in these genotypes. (Supp Fig. 2A,B)

**Figure 5:**
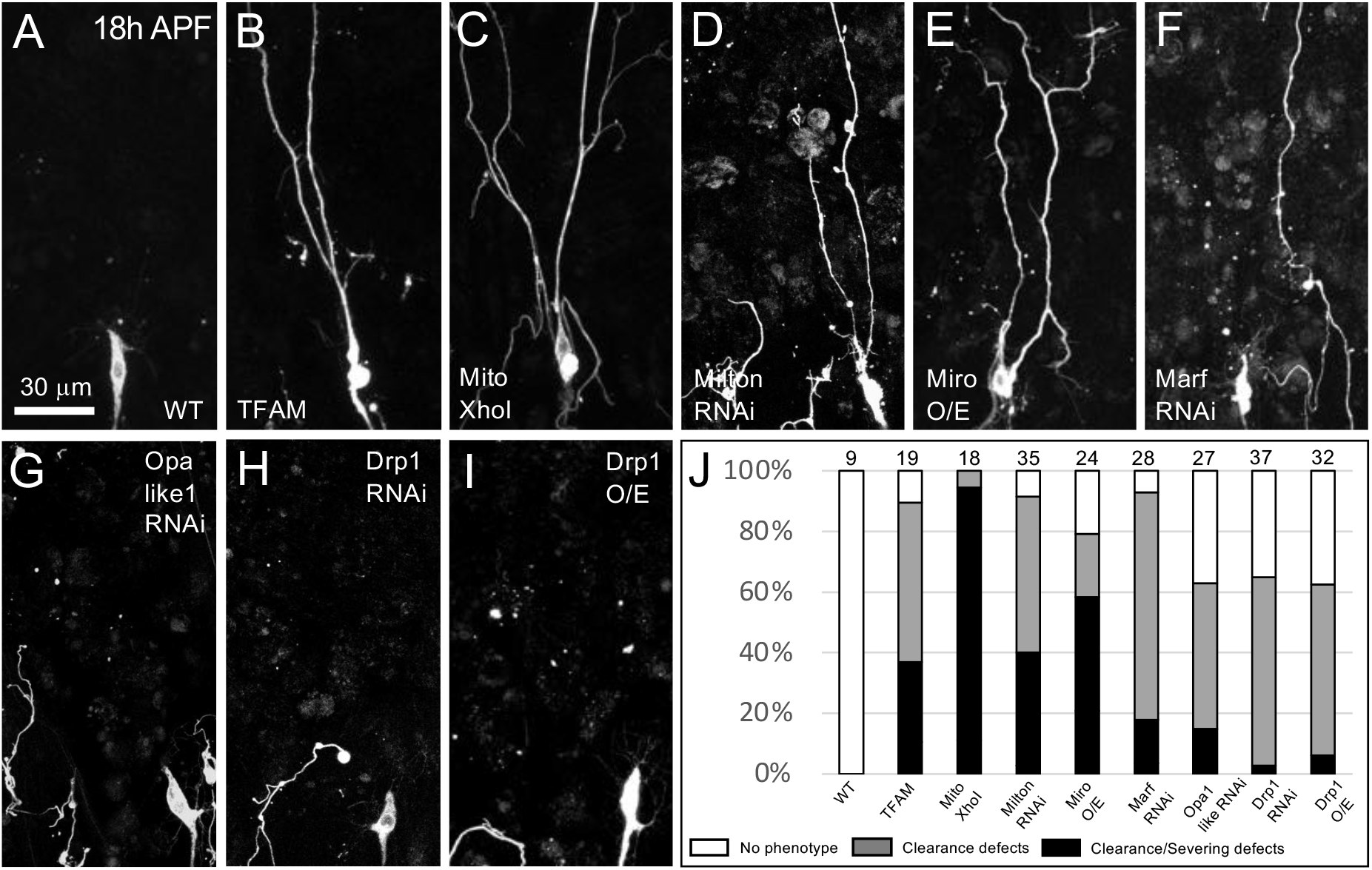
Mitochondrial function, transport and fission/fusion are important for dendrite pruning in ddaC neurons. (**A**) A wild type neuron labelled using ppkGAL4>UAS-CD8::GFP shows removal and clearance of dendrites by 18h APF. (**B,C**) Disruption of mitochondrial function blocks dendrite pruning of ddaC neurons. (**B**) Pupae expressing ppkGAL4>UAS-CD8::GFP and UAS-TFAM, a mitochondrial transcription factor, show a strong block in dendrite severing. (**C**) Disruption of mitochondrial function in neurons expressing ppkGAL4>UAS-CD8::GFP and UAS-MitoXhoI results in strong severing defects. (**D, E**) Disruption of mitochondrial transport impacts dendritic pruning of ddaC neurons. (**D**) Pupae ppkGAL4>UAS-CD8::GFP, UAS-Dicer2 and UAS-Milton RNAi, and (**E**) those expressing Miro shows defects in pruning with a robust block in severing and clearance of dendrites. ddaC neurons expressing RNAi against Marf (**F**) and Opa-like 1 (**G**) in neurons with ppkGAL4>UAS CD8::GFP and UAS Dicer2 show a block in dendrite severing and clearance. (**H**) Disruption of mitochondrial fission from expressing an RNAi against Drp1 in (**H**) or by overexpression of wild type Drp1 (**I**). Neurons expressing ppkGAL4 UAS CD8::GFP and UAS Dicer2 show disruptions in dendrite remodelling. (**J**) Chart with data from these categorised in three phenotypic groups as in Fig. 4, n numbers indicated at the top.

Mitochondrial transport is important in all cells but particularly so in neurons which have compartments distant from the cell body and have a high energy demand to fulfil (60). To determine if changing the transport and subsequent localization of mitochondria disrupts pruning, we cell-autonomously downregulated Milton, an adaptor protein, or overexpressed Miro, a critical GTPase required for transport. Both proteins have previously been shown to be necessary for mitochondrial transport along *Drosophila* motoneuron axons (61,62). Disrupting either Milton or Miro resulted in a consistent block of severing and branch clearance during dendrite pruning in ddaC neurons, similar to that seen with TFAM or MitoXhoI overexpression (Fig. 5D, E and J, Supp 1). Furthermore, we also found that this disruption of mitochondrial transport did not result in a change in Ecdysone signalling, which showed a normal onset (Fig. Supp 2 C,D).

In addition to being distributed throughout neuronal compartments via intracellular transport, the mitochondrial network is known to be highly dynamic, capable of rapid transformations in size and shape, through a balance of fission and fusion mechanisms (for review check (63)). Proper fission/fusion dynamics is also important for optimal distribution of mitochondria in the distant neuronal compartments (64,65). In majority of cell types in *Drosophila*, GTPases Marf and Opa1 are required for fusion of the outer and inner mitochondrial membrane respectively while the GTPase Drp1 regulates fission (66). Here we found that dysregulation of both fission and fusion machinery in ddaC neurons resulted in disruptions to pruning. In all cases, branch clearance was more impacted than branch severing when compared with other dysregulations of mitochondria (described above). (Fig. 5F-I and J).

To determine if these different perturbations changed the general morphology and location of the mitochondrial network within pruning neurons, we imaged GFP tagged mitochondria using Mito::GFP in individual ddaC. We found fewer mitochondria per unit length of dendrite when disrupting mitochondrial function (using MitoXhoI), as well as when perturbing mitochondrial transport (using Milton RNAi or Miro overexpression), but not when overexpressing Drp1, which was similar to wildtype neurons. (Fig. 6A-F).

**Figure 6:**
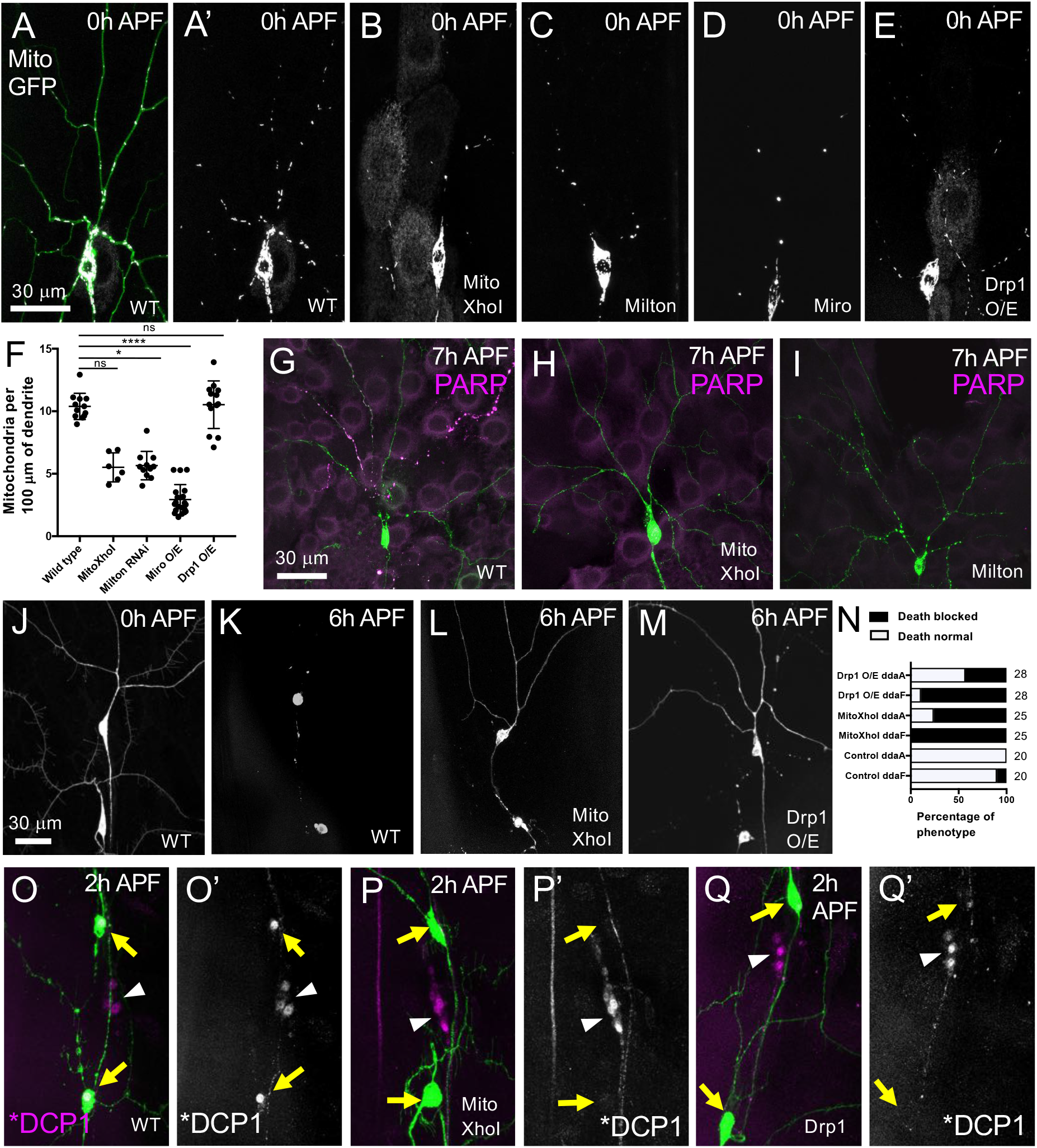
Dysregulating mitochondrial function, transport and fission/fusion changes the distribution of mitochondria and caspase activation in pruning and dying da neurons. (**A - F**) Pre-pupal neurons expressing ppkGAL4 UAS-Mito::GFP and different UAS-RNAis to perturb mitochondrial function, transport and fission. (**A**) Left panel showing a wild-type ddaC neuron expressing ppkCD4td Tomato (green) and ppkGAL4 UAS-Mito::GFP(grey), showing the distribution of mitochondria in the dendrites. (**A’**) shows the distribution of mitochondria (grey) in the dendrites, without the membrane marker. In pre-pupae expressing ppkGAL4 UAS-Mito::GFP, disruption of mitochondrial function by expression of MitoXhoI (**B**) and transport with Milton RNAi (**C**), or overexpression of Miro (**D**) decreases the number of mitochondria present in the dendrites. (**E**) In pre-pupae overexpression of Drp1 does not show a significant effect on the number of mitochondria in the dendrites. (**F**) Chart plotting the number of mitochondria per 100μm of dendrite length in genotypes shown in A. Values are reported as mean ± standard deviation and p values are reported as * for p<0.05 and ** for p <0.01. (**G - I**) Disruption of mitochondrial function and transport disrupts caspase activation in the dendrites. Wild type pre-pupae neurons expressing ppkGAL4 UAS-CD8::PARP::Venus show active cleaved-PARP in their dendrites.(**G**) The disruption of mitochondrial function by expression of MitoXhoI (**H**) or mitochondrial transport by knockdown of Milton (**I**) in ddaC neurons block caspase activation in the dendrites. (**J – N**) The disruption of mitochondrial function and fission blocks cell death in Class III neurons. (**J**) Wild type ddaF and ddaA neurons labelled using 19-12 GAL4 UAS-CD8::GFP undergo normal cell death by 6h APF (**K**). Upon disruption of mitochondrial function using MitoXhoI (**L**) and mitochondrial fission by overexpression of Drp1 (**M**) cell death is blocked in both ddaF and ddaA neurons labelled using 19-12 GAL4 UAS-CD8::GFP. The block in cell death at 6h APF is quantified and represented graphically (**N**). ddaF and ddaA neurons labelled using 19-12 GAL4 UAS-CD8::GFP, fixed and stained against GFP (green) and active DCP-1 (magenta, left panels or grey, right panels) show strong nuclear staining of DCP-1 (**O O’**) in wild type neurons at 2h APF. This active DCP-1 staining is lost when mitochondrial function is disrupted (**P P’**) or when mitochondrial fission gene Drp1 is overexpressed (**Q Q’**).

As detailed above we have found caspases to be active throughout the early stages of pruning and we next wanted to know whether they were active locally within the dendrites of neurons in which we had disrupted mitochondria function. Although SR4VH is excellent for revealing temporal and quantitative aspects of caspase activation during pruning we used mCD8::PARP::Venus here as it allows spatial visualisation of active caspases within the dendrites. Strikingly, we observed no caspase activation when we disrupted either mitochondrial physiology, with MitoXhoI, or transport, with Milton RNAi. In wild type neurons we would normally see robust PARP-IR signal in the dendrites of ddaC at 7h APF (Fig 6G-I).

As mitochondria are essential for cellular viability we checked if these disruptions impacted the survival of neurons and/or the morphology of their dendritic arborizations at larval stages and found the number of ddaC neurons remained the same, as did the numbers of primary, secondary and tertiary arbor branches (unpublished observations).

As the removal of larval neurons by apoptosis is also critical for restructuring the sensory system, we wondered whether mitochondria are also important for caspase activation and cell death and removal of class III da neurons ddaA and ddaF (Figure 6). As described, both ddaA and ddaF undergo apoptosis within 6h APF and we detect active caspases in these neurons very early during metamorphosis (see Fig 1C and Fig 3). When we disrupted mitochondrial function by expressing MitoXhoI or dysregulated fission by overexpressing Drp1, cell death was blocked (Fig 6J-N). In most cases, the cell body and some dendrites of ddaA were still visible at 6hAPF while ddaF had almost all of its dendrites as well as the cell body intact (Fig 6J-N). These changes in morphology were also mirrored in the dynamics of caspase activation. In wild type neurons at 2h APF we see that the nuclei in both ddaA and ddaF neurons stained positive for Dcp-1 (Fig 6 O, yellow arrows), which correlates with their morphology showing clear signs of cell death. We know that other dorsal neurons not labelled with the *GAL4* driver (Fig 6 O, white arrowhead) are positive for active Dcp-1. In contrast, when we imaged ddaA and ddaF neurons expressing MitoXhoI and Drp1, no active Dcp-1 was detected (Fig 6 O-Q). Thus, we show a clear link between mitochondria and caspase activation and cell death in these doomed neurons. In summary, mitochondria appear to be important players in both pruning and cell death of da sensory neurons and are important for caspase activation within these neurons during metamorphic remodelling of the sensory nervous system of *Drosophila*.

## Discussion

Nervous systems are built by both *progressive* development phenomena, such as cell division and cell growth, and *regressive* phenomena, such as cell death and pruning (67). Here we focus on the pruning and cell death of identifiable neurons in the remodelling sensory system of *Drosophila*. In our previous work we used the genetically encoded caspase probe CD8::PARP::Venus to visualise caspase activity in the dendritic branches of single pruning da sensory neurons at 6 – 7h APF (10). We found that caspase activation only occurred in dendritic branches that had been severed from the main body of the neuron and found no evidence for caspase activity early, prior to branches being cut. In contrast, our colleagues reported an early activation of caspases at 4h APF, using an antibody against cleaved human Caspase-3, and observed a suppression of branch thinning and severing in DRONC null neurons (9). As these two dataset seemed irreconcilable, we felt motivated to further investigate the timing of caspase activation and the role of other components of the apoptotic machinery during the early phases of dendrite pruning.

To look at the timing of caspase activation during pruning we first used a polyclonal antibody raised against a cleaved form of the *Drosophila* effector caspase Dcp-1. With this we found that the pruning neuron, ddaC, showed an early and weak cytoplasmic signal for ‘cleaved Dcp-1’ soon after the onset of metamorphosis, significantly earlier than our previous CD8::PARP::Venus reporter data had shown. This anti-cleaved Dcp-1 antibody recognises epitopes on cleaved Dcp-1 and on cleaved Drice (51) and so can be considered to be a good reporter of DRONC activity. These data suggests that DRONC is active early, prior to dendritic branch severing and is consistent with the data from the cleaved human Caspase-3 antibody (9).

Since in our previous work (10) we saw no ‘early’ effector caspase activity with the CD8::PARP::Venus probe, we wondered if that was due to the sensitivity of our probe or because there is no effector caspase activity at this earlier time point. An absence of effector activity was a clear possibility as a number of studies looking at non-apoptotic events, such as arista morphogenesis (52) and border cell migration (53), found that only DRONC activity is required. To address this directly we used our new genetically encoded effector caspase probe ‘SR4VH’ (40). We recently used SR4VH to describe apoptosis in newly born neurons in the ventral nerve cord during postembryonic neurogenesis (40) and found it was sensitive. The dynamics of SR4VH cleavage in the pruning neuron ddaC clearly showed, for the first time, that effector caspase activity occurs very early, prior to overt changes in the structure of proximal dendrites and much before branch severing. The improved sensitivity in revealing caspase activation could be due to a combination of features; SR4VH is a live reporter that undergoes a change in cellular localisation from the cell membrane to the nuclear compartment and the use of 4x tandem caspase cleavage sites rather than a single one. Our previous observations of active caspases in severed branches (10) had suggested to us that caspase activation was ‘held in check’ within pruning neurons because the activity was physically separated from the ‘main body’ of the neuron. These new data show that active caspases are present soon after 0h APF and are not in a physically ‘separate’ compartment. How then does ddaC deploy active caspases but not undergo programmed cell death itself? Immunostaining with the active Dcp-1 antibody pointed toward the levels of caspase activity being lower in the pruning neuron ddaC compared to the dying neurons, ddaA/ddaF. As class III (dying) and class IV (pruning) da neurons are in close proximity, on the body wall, using a live probe we could simultaneously monitor caspase activation in both. The SR4VH data mirrored observations with active-Dcp-1 antibody pointing to lower levels of caspase activity in pruning neurons than in dying neurons. A caveat to this could have been that the differences we see in the intensity of ‘cleaved’ nuclear Venus were the result of technical issues, i.e. different GAL4 levels in the two neuronal cell types or due to a ‘dilution’ of the reporter over a larger sized dendritic tree. Fortunately, having two different fluorescence proteins on either side of the caspase cleavage sites meant we could easily make a ratiometric comparison and exclude these concerns.

We know from other studies that apoptosis is not binary and that a cell may exhibit some cellular and molecular features of programmed cell death yet still survive (68). This landscape was explored by Florentin and Arama who precisely manipulated the levels of effector caspase proenzymes and showed that cellular lethality occurs once caspase activity levels reach a critical threshold (69). Below threshold, cells fail to induce apoptosis and above it a positive feedback loop accelerates the apoptotic decision (69). Ditzel et al., 2008 demonstrated how low effector caspase levels could be maintained in cells by a negative feedback regulation where DIAP inactivates caspases without degradation (70). It may be that levels of activity below such apoptotic thresholds facilitate non-lethal, non-apoptotic developmental functions, such a cell fate specification (71) and plasticity at synapses (72). This will be clearer in the future when specific targets of caspases are identified in both apoptotic versus non-apoptotic contexts.

Taken together, these data show that within the pruning neuron ddaC there is early DRONC caspase activity, that cleaves an effector caspase, either Dcp-1/Drice, and that effector caspases are active during the early phases of dendrite pruning, at levels that are clearly lower than in da neurons that are undergoing apoptosis.

Following these observations, we wondered what factors could be regulating the sub-lethal levels of caspase activation within pruning da neurons. Until now only a small number of regulators of sub-lethal non-apoptotic caspase function have been identified. Work on sensory organ precursor (SOP) development in the *Drosophila* wing imaginal discs revealed that I-kappaB kinase ɛ (IKK) indirectly regulates DRONC via phosphorylation and accelerated destruction of DIAP1, but such regulation could work independently of the RHG proteins (73). Another reported regulator of sub-lethal caspase activity is Tango7/eIF3m, which has been shown to interact with the apoptosome in testis (74), to regulate caspase activity in the salivary glands (75) and was implicated in the regulation of sub-lethal caspase activation in pruning da neurons (75). Interestingly, we found that knocking down Tango7 in ddaC neurons inhibited pruning but had no effect on caspase activation *in vivo* (Supp Fig. 2E,F).

The RHG proteins (Reaper, Grim, Hid & Sickle) are widely recognised as key regulators of cell death but have not been implicated in the regulation of sub-lethal, non-apoptotic caspase function. Our data, using MARCM clonal analysis, unequivocally shows that loss of these proteins lead to a block of dendritic pruning in the sensory neuron ddaC. To narrow down which of these proteins are required we used a series of deletions and found that mutants for both Reaper and Grim supressed dendrite pruning but neither alone was as disruptive as removing both together. Such cooperative role of RHG genes in cell death has previously been shown in the midline cells of CNS (76,77) and dMP2 neurons in late *Drosophila* embryos (78). Another possibility is that the regulatory regions of RHG genes are important for their effective function in the pruning neurons. An enhancer element located between Reaper and Grim genes called the Neuroblast Regulatory Region (NBRR) plays an important role in cell death in *Drosophila* embryonic and larval neuroblasts (79). Our mutant allele of *reaper*, just has the coding region removed, the regulatory region intact and this may result a weaker phenotype because the full cis-regulatory region allows appropriate expression of the other RHG genes (79,80). This may also explain why *H99* MARCM clones as well as *H99/XR38*, show such a strong pruning defect when compared with the single gene mutants over *H99* deficiency.

Interestingly out of the three RHG genes, we found that *hid* mutants did not result in a pruning phenotype. This was unexpected because in the context of mitochondrial caspase activation, both Reaper and Grim are known to bind to Hid to form a multimeric complex that strongly promotes apoptosis (30). Interestingly, Hid also has a wider expression pattern than Rpr or Grim and is expressed in both apoptotic as well as non-apoptotic cells (81). In case of DNA damage induced cell death like on exposure to ionising radiation, apoptosis is dependent more on Hid than on any of the other RHG genes (82). Although, it may be possible that Reaper can still localise to the mitochondria without Hid via a GH3-lipid interaction (29), multiple studies have shown Reaper to be more efficient at auto-ubiquitylating and degrading DIAP1(26,29) and more potent at inducing death (30) when present on the mitochondrial membrane. The strong block of dendrite severing in H99 MARCM single cell loss of function data we present here is striking, something we and others (83) have not seen upon removal of DRONC or following the downregulation of effector caspases (see Supp Fig 3). This difference raises the possibility that removal of the RHG proteins may result in a more significant pruning phenotype because of a failure to destroy DIAP blocking both initiator and all effector caspase function. Another possibility may be that there are as yet unknown ‘caspase independent’ functions for the RHG proteins, that are important in neuron remodelling. Abdelwahid *et al*., proposed that in addition to inhibiting DIAPs, Reaper also localises to mitochondria and permeabilises it, which is possibly a slower process than the rapid DIAP inhibition and caspase activation (32). It is possible that this slow and weak caspase activation is more significant in remodelling neurons. In addition to activation of caspases, both Reaper and Grim have also been indicated to play a role in inhibition of DIAP1 translation which, in case of Reaper, has been suggested to be crucial for cell death induction in mammalian cells (see review (84) for details). During early pupal development, DIAP1 protein levels are higher in the ddaC neurons as compared to their neighbouring ddaA/ddaF (35), suggesting that the levels of DIAP1 are differentially regulated during development. Whether the RHG proteins are involved in such mechanisms in a non-apoptotic context in these neurons is something to be investigated further.

Since the RHG proteins have been shown to intimately associate with mitochondria and because there has been an increasing body of work linking mitochondrial dynamics to caspase activation, we decided to investigate whether mitochondria are critical for remodelling of the sensory system. When we overexpressed TFAM and MitoXhoI in single neurons we saw robust disruptions in both dendrite severing and clearance. When we changed mitochondrial localization by manipulating transport and their fission/fusion dynamics in a cell-autonomous manner in ddaC neuron, we observed consistent blocks of severing and branch clearance. Notably, they also resulted in fewer mitochondria per unit length of dendrite. These mitochondrial perturbations caused a suppression of caspase activation in pruning dendritic branches. Taken together we found that the location of mitochondria within the dendritic arborizations and/or their total number is critical for normal pruning to take place.

In addition, we found that the suppression of caspase activation by mitochondrial perturbations also impacted caspases in the ‘doomed’ dying class III da neurons ddaA/ddaF and that they failed to undergo programmed cell death at the onset of metamorphosis. Although the role that mitochondria and cytochrome c play in apoptosis in *Drosophila* has been an open and somewhat awkward question (85), our data here brings insight to and strong support for the idea that mitochondria are playing a key role in caspase activation *in vivo* in dying *Drosophila* sensory neurons.

In summary, our study shows that during da neuron pruning, caspases are active earlier than previously thought and that effector caspase activity is lower in pruning than in dying neurons. We reveal that the pro-apoptotic factors Reaper and Grim are required for neuronal pruning in the sensory system of *Drosophila*. We find that the location and/or function of mitochondria are critical for pruning and caspase activation in both remodelling and dying neurons. These data on the sub-lethal regulation of caspases are consistent with a growing body of work in flies of an axis of mitochondrial fission/fusion and caspase activation. Looking forwards, we hope that by genetically tagging these RHG proteins and comparing their dynamics and localisation in pruning versus dying neurons will give us greater insights into their mechanism of action in sub-lethal, non-apoptotic processes.

## Author contributions

AM and DW conceptualised the project, designed experiments and wrote the manuscript. Experiments and analysis were performed by AM except for Figure 2 and the intensity analysis in Figure 3, which were done by SP. Hid and Reaper mutants were generated by SK. All authors read the final manuscript and provided feedback.

## Acknowledgements

We would like to thank Nicolas Loncle for initial observations on caspase activation in sensory neurons with *Apoliner* and SR4VH. We would like to thank Bloomington Stock Centre, Tadashi Uemura, Kristin White, Joe Bateman and Alex Whitworth for providing several fly lines. Thanks to FengWei Yu for providing us with the antibody against Sox 14. Thanks to Paul Conduit for his generosity in providing reagents, imaging facility and time to generate some key data for the manuscript. Many thanks to Joe Bateman and Alberto Baena-Lopez for reading and feedback on the manuscript. All the Williams lab members for their help and assistance with maintaining and sending fly stocks.

---

**Supplemental Figure 1:**
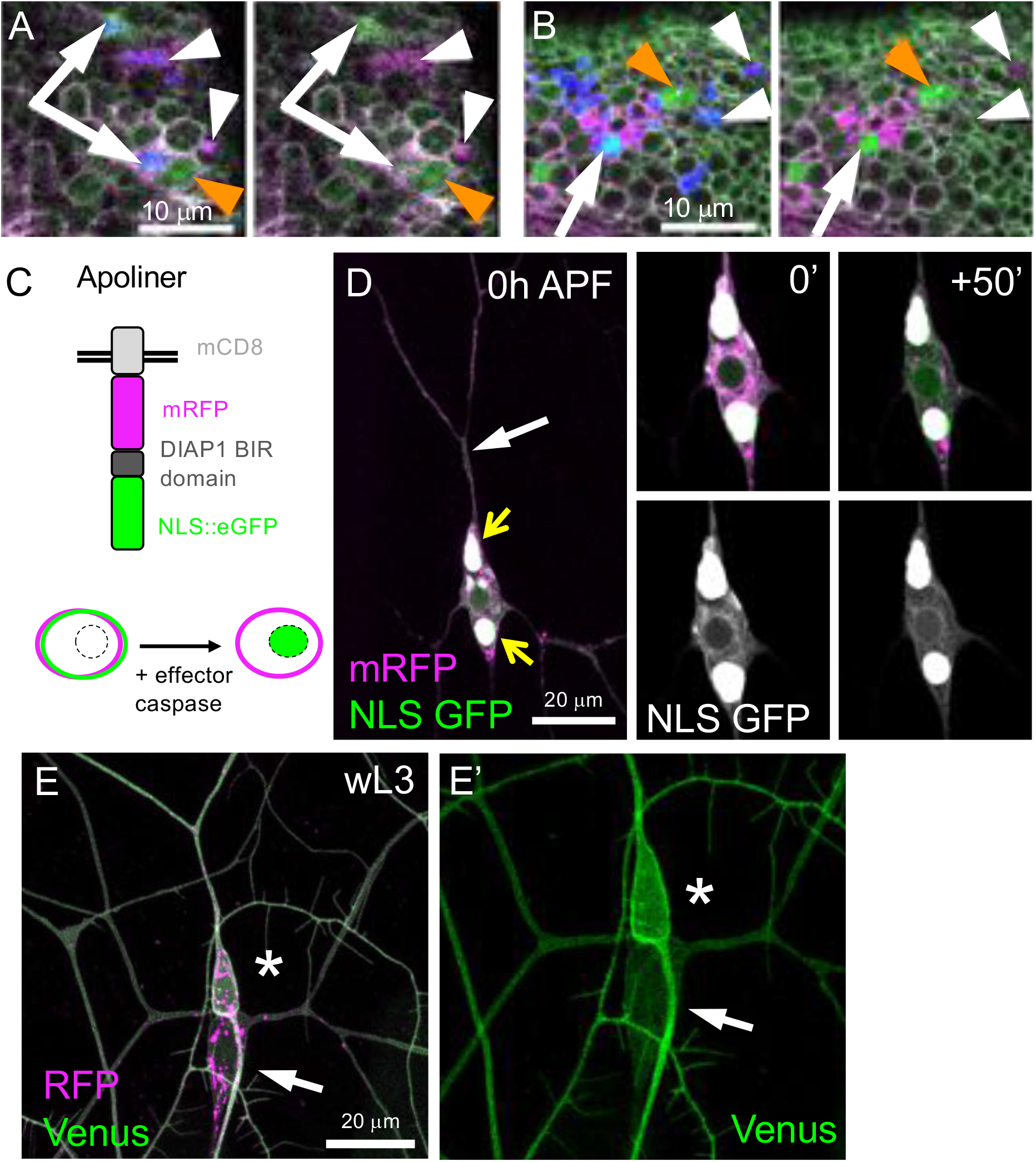
**(A)** Wing discs of Control females and (**B)** wing discs of *hs-hid* males exposed to 1h heat-shock at 37°C reveal cells at successive stages of cell death. SR4VH reporter expression (magenta = RFP; green = Venus) together with immunolabelling for the cleaved effector caspase Dcp-1 (blue) orange arrowheads = nuclear Venus without cleaved Dcp-1 (early-stage); white arrows = both nuclear Venus and cleaved Dcp-1 (mid-stage); and white arrowheads = pyknotic cells/dead cell membranes with RFP and cleaved Dcp-1 (late-stage); Scale bars = 10 μm. (**C**) Schematic representation of genetically encoded Apoliner probe and its mechanism of action. (**D**) Left panel shows a prepupa with ddaC expressing UAS-Apoliner. Sensory neurons showed large accumulations of the probe within the Golgi (yellow arrows). The first and last time points from a movie showing a single slice through the middle of the soma and the weak accumulation of GFP (green, top panel and grey scale, bottom panel) in the nuclear region. The dendrites are still attached to the cell body (**D**, white arrow), suggesting weak active caspases in ddaC. (**E**) 2x ppkGAL4 expressing UAS-SR4VH in wandering third instar larva (wL3) Some RFP accumulates in vesicles but the membranes of both class III and class IV neurons are evenly labelled. No nuclear GFP is found in any of these sensory neurons during this period. (**E’**) Shows Venus channel. The image is magnified to focus on the cell bodies.

**Supplemental Figure 2:**
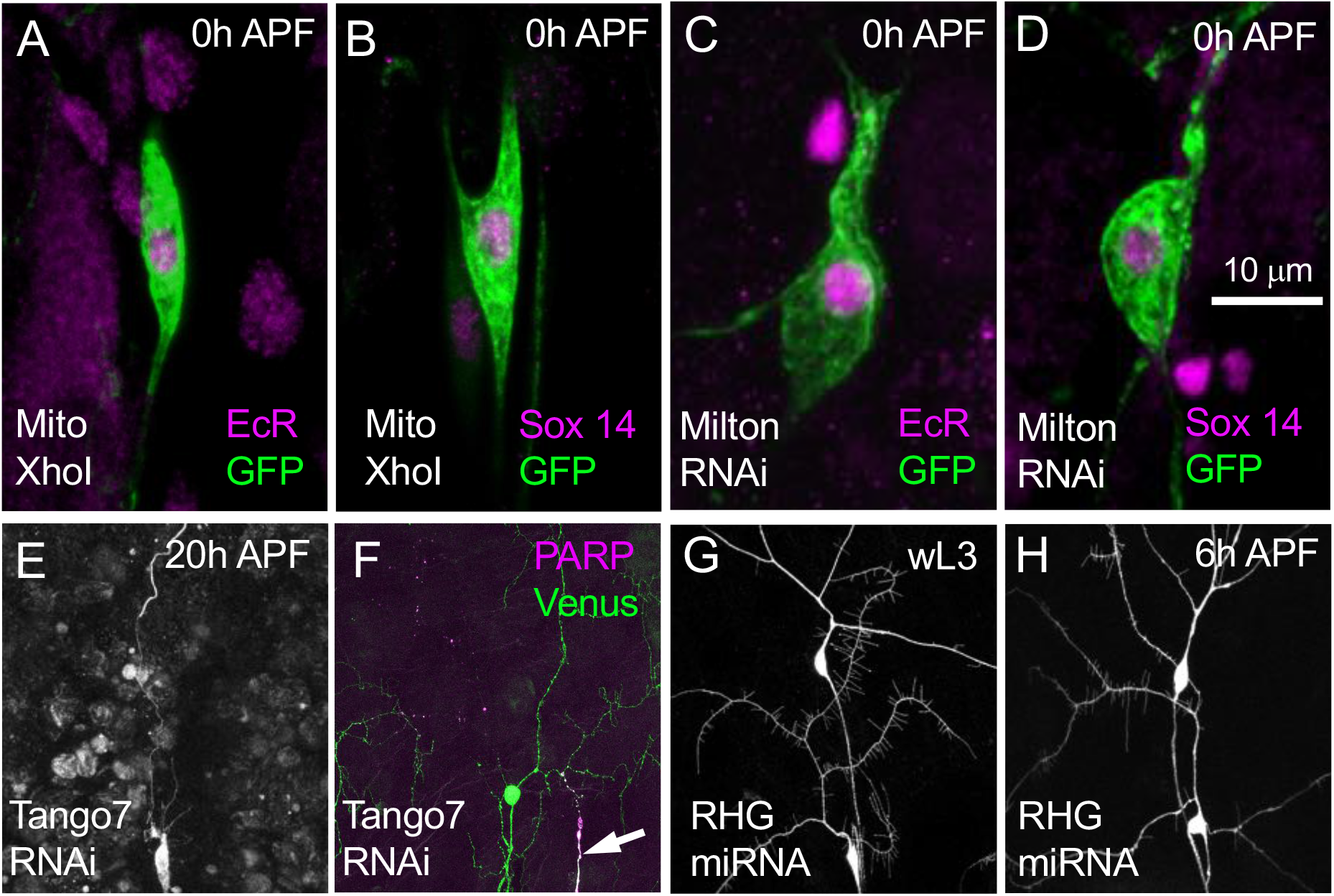
(**A,B**) Pre-Pupae expressing MitoXhoI in ppkGAL4 UAS CD8::GFP neurons fixed and stained for EcR (**A**, magenta) and GFP (green) or Sox14 (**B**, magenta) and GFP (green). These show normal expression of both hormonally gated transcription factors. (**C,D**) Pre-Pupae expressing Milton RNAi in ppkGAL4 UAS-CD8::GFP neurons fixed and stained for EcR (**C**, magenta) and GFP (green) or Sox14 (**D**, magenta) and GFP (green). (**E)** ddaC neurons labelled with ppkGAL4 UAS CD8::GFP at 20h APF showing pruning defects when Tango7 is knocked down using UAS-Tango7 RNAi **(F)** A 7h APF pre-pupae expressing ppkGAL4 UAS CD8::PARP::Venus and UAS Tango7 RNAi. This was immunostained against anti-GFP (green) and anti-PARP (magenta) and showed detectable levels of cleaved PARP (arrows) within the dendrites of the ddaC neurons. **(G,H).** The disruption of RHG proteins in Class III neurons. (**G**) ddaF and ddaA neurons expressing RHG-RNAi in in 3^rd^ instar expressing in larva look normal. **(H)** ddaF and ddaA neurons expressing RHG-RNAi do not show signs of apoptosis at 6h APF.

**Supplemental Figure 3:**
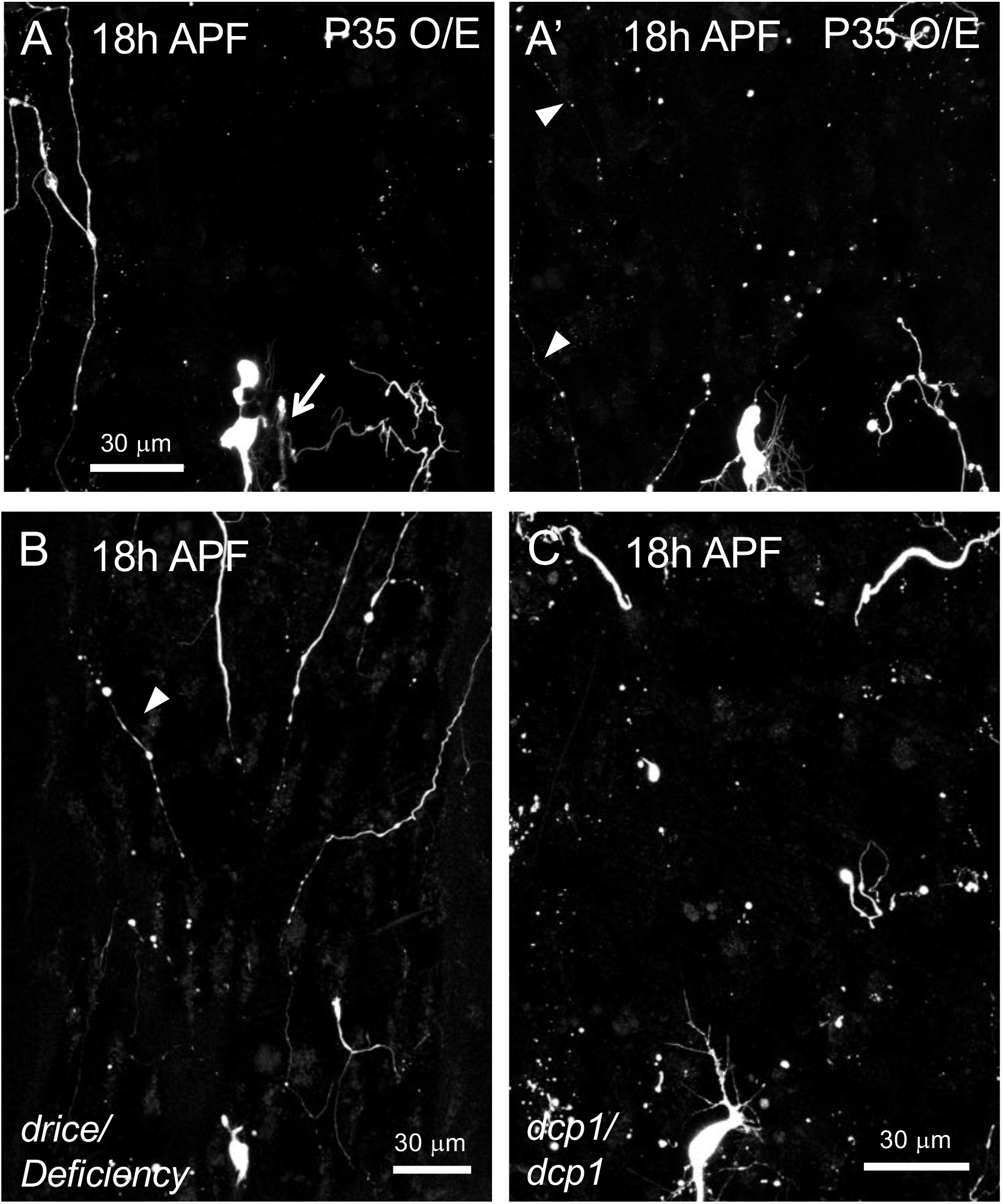
**(A-A’)** ddaC neurons labelled using ppkGAL4>UAS-CD8::GFP, UAS-p35 at 18h APF expressing two copies of the baculovirus protein P35. These show pruning defects ranging from **severing (A, arrow)** to clearance defects **(A’, arrowheads)**. **(B)** ddaC neurons labelled using ppk-eGFP showing clearance defects in Drice mutant over Drice deficiency at 18h APF. **(C)** ddaC neurons labelled using ppkeGFP showing clearance defects in DCP-1 homozygous mutants at 18h APF.

**Supplementary Movie 1: Time-lapse of dorsal multiple dendrite neuron (dmd1) expressing SR4VH reporter in prepupa.** Time-lapse movie of dmd1 neuron (arrow) in a prepupa, 2h after puparium formation. Sensory neurons expressing SR4VH reporter under the control of elav^C155^ GAL4 imaged every 10 minutes. GFP leaves the membrane and accumulates in the nucleus over the time course revealing caspase activation. Other sensory neurons in close proximity already showing robust caspase activation.

**Supplementary Movie 2: Time-lapse movie of a class IV da neuron ddaC and class III da neuron ddaA expressing SR4VH reporter in prepupa.** Time-lapse movie of a ddaC neuron (magenta arrow) and a ddaA neuron (green arrow) starting at white prepupa (0h APF), imaged every 5 minutes. Sensory neurons expressing SR4VH reporter under the control of two copies of ppk-GAL4. Timelapse reveals caspase activation by the accumulation of GFP in the nucleus of ddaC (pruning, class IV da neuron) and a robust accumulation of GFP in the nucleus of ddaA neuron (dying, class III da neuron) over the time course.

